# Dopamine shapes brain metastate dynamics

**DOI:** 10.64898/2026.01.30.702810

**Authors:** Daisuke Matsuyoshi, Yasuyuki Kimura, Keisuke Takahata, Yoko Ikoma, Chie Seki, Ming-Rong Zhang, Makoto Higuchi, Tetsuya Suhara, Makiko Yamada

## Abstract

Dopamine’s influence on large-scale network dynamics, especially on the default mode network (DMN), remains uncertain, as fMRI studies have produced mixed results. One likely contributor to these discrepancies is reliance on traditional functional connectivity analyses, which typically derive a single metric (e.g., the Pearson correlation coefficient) from the entire time series and thus fail to capture network dynamics. To address this issue, we combined a dopaminergic challenge (mazindol, a dopamine transporter [DAT] reuptake inhibitor), PET, resting-state fMRI, and hidden Markov modeling (HMM) to characterize time-varying alterations in human large-scale functional networks following acute DAT blockade. We found that mazindol-induced increases in endogenous dopamine altered the balance between the brain’s functional “metastates,” two recurrent higher-order network configurations that each encompass multiple HMM-derived brain states. Mazindol increased the time participants spent in an internally oriented cognitive metastate and decreased the time spent in a sensorimotor–perceptual metastate, with the DMN showing the most pronounced lengthening. In exploratory analyses, declines in [¹¹C]raclopride binding, a PET index of D2 dopamine receptor availability reflecting increased striatal extracellular dopamine levels, tended to show a positive correlation with the prolongation of these cognitive states. These findings indicate that dopamine is closely linked to shifts from sensorimotor and perceptual to cognitive brain metastates, potentially underpinning the prioritization of internally oriented over externally driven psychological processes. Our results highlight the importance of dynamic, time-resolved connectivity approaches for understanding neuromodulatory actions in the human brain and suggest that dopamine helps regulate the dynamic balance between functionally competing large-scale brain networks.

## 1. Introduction

A prevailing view in neuroscience posits that complex cognition and behavior arise not from the isolated work of individual brain regions, but from the coordinated interplay of distributed brain networks [1–3]. These large-scale functional architectures, such as the default mode, salience, and executive control networks, must flexibly reconfigure in response to ever-changing internal and external demands, thus enabling goal-directed behavior [4–6]. The orchestration of these rapid network reconfigurations critically depends on neuromodulatory systems. By releasing neurotransmitters such as dopamine, acetylcholine, and serotonin, these systems not only transmit information but also fundamentally alter the biophysical properties of neurons and synapses, thereby sculpting the brain’s entire functional landscape [7–9]. Dysfunction in these systems is implicated in a range of neuropsychiatric conditions [10–12].

Among these neuromodulators, dopamine plays a central role in motor control, reinforcement learning, and higher-order cognition [13–16]. It is thought to act as a master regulator of network topology by modulating the signal-to-noise ratio of neural firing [17–19], stabilizing network dynamics [18, 20–23], and maintaining the functional specialization of distinct brain systems [24–27]. Recent work has demonstrated that the distribution of dopamine receptors aligns with the principal gradient of cortical functional organization [28], suggesting a foundational role for dopamine in shaping the brain’s unimodal-to-transmodal hierarchical differentiation. These findings underscore dopamine not merely as a neurochemical messenger, but as a key architect of the brain’s functional organization. Understanding such global neuroregulatory roles of dopamine is essential for linking synaptic neurochemistry to brain dynamics and, ultimately, to cognition and behavior.

The default mode network (DMN) is one of the most robustly identified large-scale brain networks. It was originally described as a set of cortical regions that deactivate during externally focused, attention-demanding tasks but show heightened activity during quiet, wakeful rest [29]. A vast body of research has since established the DMN as a principal neural substrate for internally generated cognition [30, 31]. It plays a central role in a suite of self-referential processes, including autobiographical memory retrieval [32–34], prospective simulation [35–37], social cognition [38–40], and mind-wandering [41–45].

The DMN is anatomically and functionally distinct from early sensory and motor cortices, which subserve exogenously driven sensory processing and motor control [46–48]. This dissociation suggests a fundamental organizing principle of the brain, reflecting a dynamic competition between internally oriented and externally oriented processing. The DMN is therefore not a passive “idling” state but an active system that integrates information drawn from autobiographical, prospective, and social-cognitive domains to construct a coherent “internal narrative” and sustain a continuous sense of self [31]. It has been suggested that the DMN may implement these functions via “offline” computations, including internal model maintenance and optimization [49, 50].

Despite the importance of both dopaminergic systems and the DMN, their interaction remains poorly understood. Existing human studies have yielded conflicting results: some report that dopaminergic modulation *decreases* DMN activity and connectivity [51, 52], others report *increases* [53, 54], and still others find *no significant change* [55, 56]. By contrast, studies in clinical populations, including ADHD and Parkinson’s disease, have consistently shown that dopaminergic medication *normalizes* aberrant DMN connectivity [57–60]. For instance, Zhong et al. [60] reported that unmedicated patients with Parkinson’s disease showed atypical decreases in DMN connectivity compared to healthy controls, which was largely normalized following levodopa administration. Thus, dopaminergic effects on the DMN appear more robust in pathology than in health. This contrast suggests that an important moderating factor may be missing from current research paradigms.

We propose that one major source of this inconsistency is methodological rather than biological, arising from an overreliance on static functional connectivity (sFC) analysis. sFC quantifies connectivity by calculating a single metric (e.g., a Pearson correlation coefficient) across an entire fMRI time series, implicitly assuming that network relationships remain stable over time (Figure 1). However, the brain exhibits rapid transitions between distinct, quasi-stable patterns of network co-activation, commonly described as “brain states,” which can be further grouped into higher-order “brain metastates” to form a temporal hierarchy of network configurations that the brain tends to cycle through [5, 61]. Time-averaging across these distinct states is analogous to taking a long-exposure photograph, which integrates a moving scene into a single static image; the resulting composite obscures the unique features of individual brain states. We hypothesized that dopamine may not simply increase or decrease specific connectivity, but rather alter the *temporal dynamics* of these states—changes that typical sFC is blind to. This perspective has motivated a shift toward dynamic functional connectivity (dFC) analyses, which capture transient fluctuations in functional coupling and have proven to be more sensitive markers of brain function and behavior than traditional sFC [62–65]. Resolving the dopamine-DMN controversy will likely require moving beyond static snapshots and characterizing time-resolved patterns of network organization, sometimes referred to as the “chronnectome” [66].

**Figure 1.**
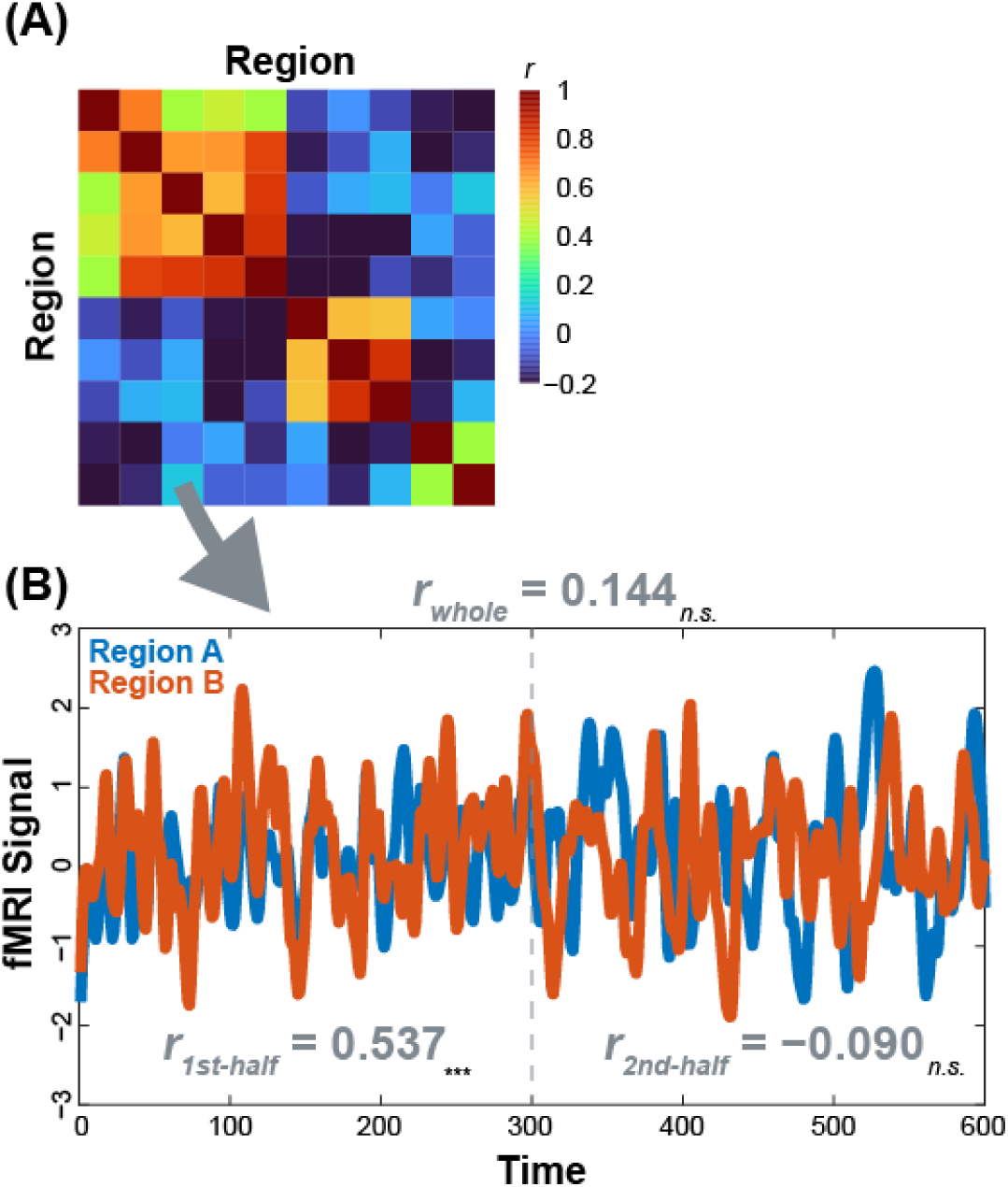
Static functional connectivity analysis does not capture time-varying connectivity dynamics. (A) Correlation matrix heatmap as seen in a typical static functional connectivity analysis. (B) Example BOLD time series from a correlation-matrix entry. Although a conventional static analysis spanning the entire time series classifies its functional connectivity as “nonsignificant” (*r* = 0.144), split-half correlations diverge: the significant first half (*r* = 0.537) and nonsignificant second half (*r* = −0.090), revealing temporal dynamics that a single static estimate obscures. This example clearly illustrates a core limitation of static functional connectivity analysis and highlights the importance of dynamic functional connectivity analysis.

Based on this dynamic framework, we reasoned that dopamine’s fundamental role is not to alter static connectivity patterns but to modulate the temporal organization of brain states. To test this hypothesis, we employed a multimodal approach that combined pharmacological intervention (mazindol, a dopamine transporter [DAT] reuptake inhibitor) using [^11^C]raclopride and [^18^F]FE-PE2I positron emission tomography (PET) and resting-state functional MRI (rfMRI). Oral administration of mazindol blocks presynaptic DAT, thereby increasing extracellular dopamine in the striatum. The resulting competition with [^11^C]raclopride at D2/3 receptors reduces binding potential (*BP*_ND_); this reduction serves as an in vivo index of dopamine release for each individual, as shown in our previous study [67]. In parallel, [^18^F]FE-PE2I PET quantifies baseline DAT availability and its occupancy under mazindol administration, enabling us to estimate how strongly the drug engages presynaptic transporters in each participant. Taken together, the raclopride- and FE-PE2I-based measures provide complementary information on both the magnitude of dopamine release and the degree of pharmacological DAT blockade across individuals.

Critically, our analytical framework overcomes the limitations of sFC approaches by employing Hidden Markov Models (HMMs) [5]. HMMs are a dFC method that captures recurring latent brain states and their temporal dynamics, such as fractional occupancy (FO, the proportion of time participants spent in each state) and transition probability (TP), with high temporal precision. Unlike sliding-window or time-frequency dFC approaches, HMMs avoid arbitrary windowing and infer discrete latent states on a timepoint-by-timepoint basis (one TR), allowing the detection of brief and frequent reconfigurations that windowed methods smear. HMMs are thus well-suited to quantify dopamine’s influence on temporal network dynamics.

Our central prediction was grounded in evidence that dopamine supports higher-order cognition by stabilizing network dynamics [20–22], including those underlying working memory. In a dynamic context, such a stabilization would be expected to prioritize internal, goal-relevant cognitive processes over competing sensory inputs [23, 68]. Given the well-established functional opposition between the DMN (supporting internal cognition) and sensorimotor and perceptual networks [5, 46–48], we hypothesized that increased dopamine release would shift the balance between these functionally opposing states, leading to a temporal reorganization of neural resources. Specifically, we predicted that mazindol-induced increases in endogenous dopamine would stabilize the DMN-led cognitive brain metastate at the expense of the sensorimotor-perceptual metastate, manifesting as a prolongation of time spent in the cognitive metastate and a corresponding reduction in the sensorimotor-perceptual metastate. To further probe this proposed mechanism, we also explored whether the magnitude of this dynamic reallocation would correlate with individual differences in striatal dopamine levels as measured by PET. We focused on the dopamine-induced prolongation of the cognitive brain metastate because the striatum is a key hub where dopamine integrates motivational signals to modulate cognitive stability and flexibility [69–71]. Demonstrating such a covariation would provide additional support for a link between molecular neurochemistry and the macroscale brain dynamics that govern the prioritization of self-generated over stimulus-driven psychological processes.

## 2. Materials and Methods

### Ethics Statement

The study was approved by the Ethics and Radiation Safety Committee of the National Institute of Radiologic Sciences (currently the National Institutes for Quantum Science and Technology), Chiba, Japan, and was performed in accordance with the ethical standards laid down in the 1964 Declaration of Helsinki and its later amendments. All participants provided written informed consent prior to participation. The study was registered with the University Hospital Medical Information Network Clinical Trials Registry (UMIN000008232). We also used data from the Human Connectome Project (HCP) datasets, which were acquired, shared, and approved by the Institutional Review Board of Washington University in St. Louis. Informed consent was obtained for each participant, and all open-access data were de-identified.

### Participants

Sixteen healthy young adults participated (16 males; mean age: 22.4 years; range: 21-25 years). All had normal or corrected-to-normal vision, and none reported a history of neuropsychiatric disorders.

### Pharmacological intervention and PET/MRI data acquisition

The participants underwent one MRI scan and two PET scans after receiving either mazindol or a placebo orally on two different days. On the first day, half of the participants received mazindol, and on the second day, they were given a placebo. The order was reversed for the remaining participants. The mean interval between the two experiments was 25.4 days (range 14-56 days). On each day, participants were administered 1.5 mg mazindol (milled tablets blended with 0.5 g lactose hydrate extra fine crystal; Sanorex® Tablets [Novartis Pharma AG, Basel, Switzerland] and Lactose “Hoei” Extra Fine Crystal [Pfizer, New York, USA]), a DAT reuptake inhibitor, or placebo (0.5 g lactose hydrate extra fine crystal) orally at around 10:30 A.M.

We acquired MR images using a Siemens Verio 3T scanner (Siemens, Erlangen, Germany) equipped with a 32-channel head coil, 1.5 hours after the drug administration. We collected anatomical 3D T1-weighted (T1w) images (Magnetization-Prepared Rapid-Acquisition Gradient-Echo sequence, repetition time [TR] = 2.3 s; echo time [TE] = 1.95 ms; flip angle = 9°; 0.49 mm × 0.49 mm in-plane resolution; slice thickness = 1 mm; 176 sagittal images) and used a single-band, gradient-echo echo-planar imaging (EPI) pulse sequence to collect T2*-weighted images during resting (TR = 2.0 s, TE = 25 ms; flip angle = 90°; 3.75 mm × 3.75 mm in-plane resolution; slice thickness = 3.8 mm; 38 axial slices; 204 volumes). Participants were instructed to open their eyes and fixate on a central cross during the rfMRI scan. Participants viewed stimuli on a 23-inch LCD monitor (BOLDScreen, Cambridge Research Systems, Rochester, United Kingdom) through a mirror mounted on a participant’s head coil.

We injected [^11^C]raclopride 3 hours and [^18^F]FE-PE2I 5 hours after the administration of mazindol or placebo. Mean injected does were approximately 219 ± 19 MBq [1.7 ± 2 nmol] for placebo and 230 ± 16 MBq [1.2 ± 0.4 nmol] for mazindol for [^11^C]raclopride, and 187 ± 21 MBq [0.6 ± 0.3 nmol] for placebo and 179 ± 12 MBq [0.7 ± 0.6 nmol] for mazindol for [^18^F]FE-PE2I. [^11^C]raclopride indexes dopamine D2/3 receptor availability, whereas [^18^F]FE-PE2I selectively labels DAT, reflecting presynaptic terminal integrity and reuptake capacity. This dual-tracer approach provides complementary receptor- and transporter-level measures of dopaminergic function in vivo. We used a PET camera (SET-3000GCT/X, Shimadzu, Kyoto, Japan) to acquire 3D dynamic PET images. For [^11^C]raclopride, we took 38 frames with increasing duration from 30 seconds to 5 minutes over 60 minutes. For [^18^F]FE-PE2I, we took 35 frames over 90 minutes. We corrected PET images for attenuation, scatter, and head movement for the late frames of the PET images using PMOD 3.7 (PMOD Technologies, Zürich, Switzerland).

Dynamic PET data were quantified voxel-wise using the multilinear reference tissue model (MRTM2) with cerebellar cortex as the reference region [67]. This yielded parametric images of *BP*_ND_ for each tracer. For each participant and condition (placebo vs. mazindol), *BP*_ND_ maps were coregistered to each subject’s T1w image, then spatially normalized (MNI152). We used the Oxford-GSK-Imanova Structural–anatomical Striatal Atlas to define our striatum ROI [72]. Percent change/decrease in *BP*_ND_ was computed as: %decrease = ((*BP*_placebo_ − *BP*_mazindol_) / *BP*_placebo_) × 100. For [^11^C]raclopride, this percentage reflects task-free changes in extracellular dopamine (a greater dopamine concentration results in a greater percentage decrease). For [^18^F]FE-PE2I, the same calculation indexes DAT occupancy by mazindol.

### MRI preprocessing

We modified the Human Connectome Project’s minimal processing pipelines [73, 74] to preprocess our single-band fMRI data, as single-band fMRI data can also significantly benefit from HCP-style preprocessing [75]. The modifications included skipping of myelin (T1w/T2w) map-based B_1_-bias correction and image-distortion correction using a field map.

Our preprocessing pipeline (Fig. 2) included slice timing correction, gradient distortion correction, motion correction, averaging of day 1 and day 2 distortion-corrected T1w images, fMRI to T1w registration using FreeSurfer’s BBRegister [76], MNI registration, intensity normalization, and brain masking for volume-based preprocessing. Surface-based preprocessing comprised parcel-constrained resampling from individual to atlas subcortical voxels (2 mm FWHM), cortical ribbon-based volume to surface mapping with exclusion of noisy voxels onto native mesh, surface resampling to a registered 32k mesh, 2 mm FWHM Gaussian surface smoothing, ICA-based automatic denoising using ICAFIX [77, 78] with high pass filtering at 0.01 Hz, and multi-modal surface matching (MSM) with group template maps [79, 80]. We did not perform global signal regression. The result is an accurately coregistered dense time series of 91,282 grayordinates. For a conventional sFC analysis, cortical parcellation was performed using FreeSurfer’s aparc.a2009s atlas [81], which divides the cerebral cortex into 148 anatomically defined regions. We further selected 30 DMN-related parcels from the atlas (15 from each hemisphere; G_and_S_transv_frontopol, G_and_S_cingul-Ant, G_and_S_cingul-Mid-Ant, G_and_S_cingul-Mid-Post, G_cingul-Post-dorsal, G_cingul-Post-ventral, G_front_sup, G_pariet_inf-Angular, G_precuneus, G_temporal_inf, S_interm_prim-Jensen, S_pericallosal, S_subparietal, S_temporal_inf, and S_temporal_sup) to increase statistical sensitivity. We thus computed parcel-wise sFC matrices at two scales and tested for differences between conditions.

**Figure 2.**
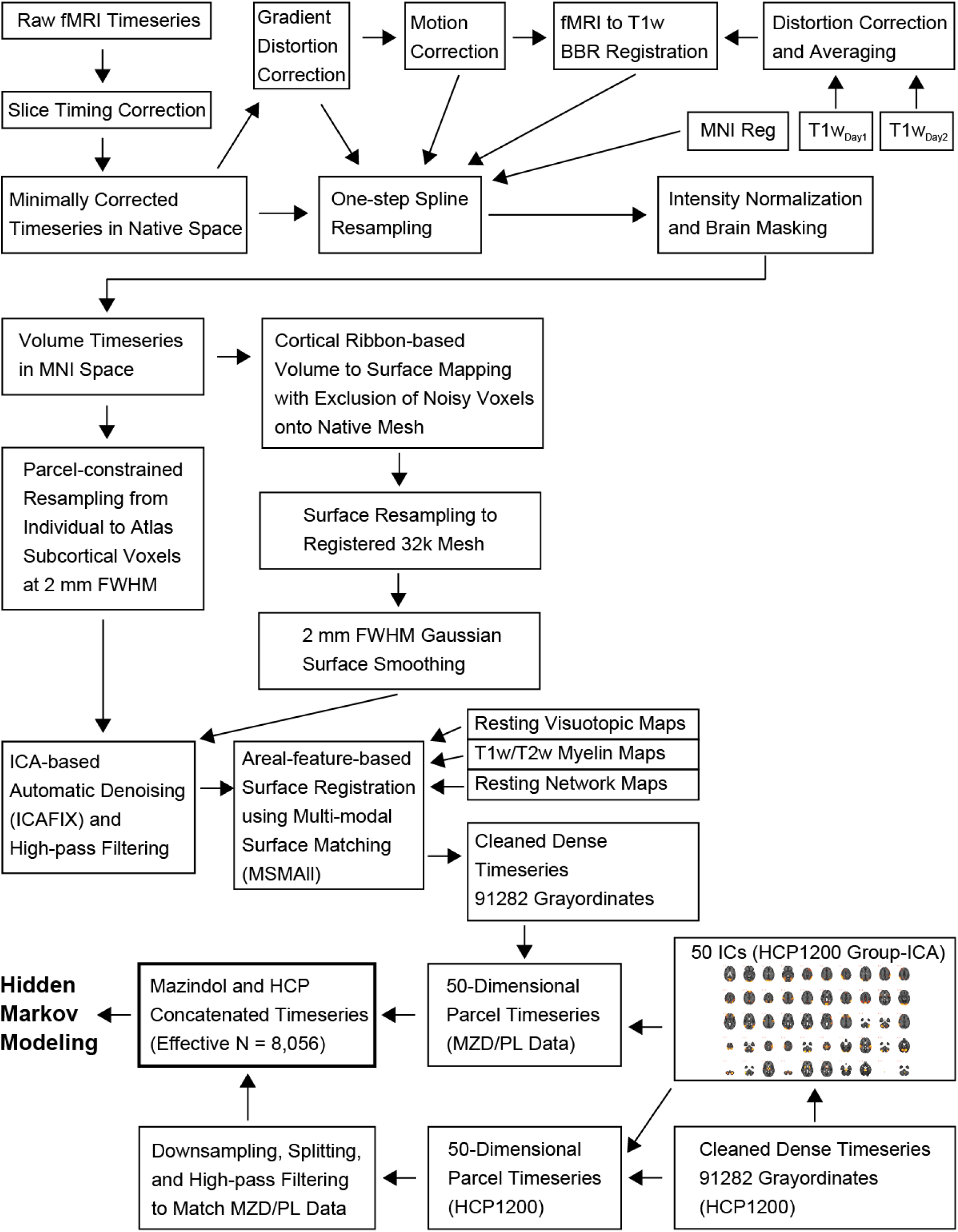
MRI preprocessing pipeline.

### HMM on rfMRI

To examine time-varying brain dynamics, we applied HMM [5] to our rfMRI data. HMM models the observed BOLD signal as arising from a Markov process that switches among unobservable “hidden” states, termed brain states, which correspond to recurrent configurations of large-scale brain networks. This approach enabled us to infer a timepoint-by-timepoint sequence of brain states and to quantify time-resolved state dynamics using FO, transition probabilities, and switching rate.

To improve the stability of state estimates by increasing the total number of observations [82], we combined our data with the WU-Minn HCP 1200 Subjects Data Release (hereafter, HCP1200), which includes complete T1w and rfMRI datasets from 1,003 participants. To align the spatial domains between HCP1200 and our datasets, both datasets were parcellated into 50 spatial components using group spatial independent component analysis (group ICA) applied to the HCP1200 data. We then performed four temporal preprocessing steps to align the two datasets’ temporal domains. First, the 50-dimensional HCP1200 data were down-sampled to a virtual TR of 2 s, reducing the original 1,200 volumes/run (TR = 0.72 s) to 432 volumes/run (virtual TR = 2 s). Second, to maximize use of the HCP1200 dataset, we split each of the four original runs into two segments, yielding eight virtual runs in total. This resulted in an effective sample size of N = 8,024 (1,003 × 8) for our data analysis, considering the scan length. Third, we discarded the initial 15 volumes/timepoints from both datasets to ensure the signal had reached equilibrium, and high-pass filtered the split HCP1200 data at 0.01 Hz. Thus, we used 189 + 189 volumes from the down-sampled 432 volumes/run of the HCP1200 data (discarded the initial 15 volumes, used 189, used 189, and discarded the last 39 volumes/timepoints). Finally, we merged the processed HCP1200 data with our data. The resulting temporally concatenated time series (effective number of participants [scans] × number of time points × number of components = 8,056 [1,003 × 8 + 16 +16] × 189 × 50) were standardized so that each scan (run), subject, and component has a mean of 0 and SD of 1. To further reduce noise, we performed principal component analysis on the temporally concatenated, standardized time series, retaining components that together explained 90% of the variance.

We modeled the final fMRI time series with an HMM using the HMM-MAR toolbox, in which each state is represented by a multivariate Gaussian with a state-specific mean and covariance. Following Vidaurre et al. [5], who used the same HCP dataset, we set the number of states in the HMM as 12. The model was fit at the group level on temporally concatenated data, yielding subject- and condition-specific state-probability time courses and a TP matrix specifying the probability of moving from any state *i* to state *j*. FO for each state was defined as the proportion of time a participant spent in that state, computed as the sum of posterior probabilities of that state across time divided by the number of time points. Switching rate was defined as the number of transitions between distinct states per TR, computed from the most-probable state sequence decoded from the posterior time courses (Viterbi path).

To identify hierarchically organized metastates, defined as superordinate cyclically recurring sets of brain states [5], we first applied modularity-maximization-based clustering to the TP matrix, treating it as a directed, weighted graph and using the Leiden algorithm with 1,024 iterations [83]. Leiden partitions the nodes (brain states) into communities (metastates) by maximizing within-community connectivity relative to between-community connectivity, and unlike Louvain, it guarantees well-connected communities. Second, we constructed the FO Pearson correlation matrix to capture how time spent in each state covaried across individuals. We then applied hierarchical clustering using the unweighted pair group method with arithmetic mean (UPGMA) to characterize the hierarchical organization of brain states.

### Integration of PET and rfMRI HMM

To test whether individual differences in striatal dopamine release covaried with mazindol-induced changes in brain-state dynamics, we correlated PET-derived dopamine indices with HMM-derived FO across individuals. Dopamine release was indexed by the percent decrease in [^11^C]raclopride *BP*_ND_ within the striatal ROI (mazindol relative to placebo). As a supplemental sensitivity check, we also quantified mazindol DAT occupancy (target engagement at DAT) as the percent decrease in [^18^F]FE-PE2I *BP*_ND_ within the same ROI. For each HMM state, drug effects on temporal dynamics were summarized as ΔFO = FO_mazindol_ − FO_placebo_, and correlation coefficients were computed between PET indices and ΔFO across individuals.

### Statistical analysis

All statistical tests used an *α*-level of 0.05. For sFC, we controlled for multiple comparisons across all possible connections using the Benjamini–Hochberg false discovery rate (FDR) procedure (*q* < 0.05). For state-wise comparisons, we applied the same FDR procedure within each metastate-defined family of states. Between-variable associations were quantified using Pearson’s product–moment correlation coefficient; Kendall’s *τ* was additionally computed as a rank-based sensitivity analysis.

We analyzed FO and TP while accounting for the compositional constraints inherent to both metrics. For FO, fractional occupancy is compositional across the set of metastates identified by the HMM; within each condition, occupancies sum to one 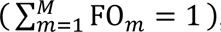, where *M* denotes the number of metastates. For TP, transition probabilities are compositional within each source metastate; for a given source metastate *i*, the probabilities of transitioning to all destination metastates sum to one 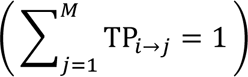. In the special case of two metastates (*M* = 2), this reduces to FO_metastate1_ + FO_metastate2_ = 1 and TP*_i_*_→_*_i_* + TP*_i_* _→_*_j_* = 1, where *j* denotes the other metastate. Accordingly, in the two-metastate case, the analysis focuses on the own-state TP for each source metastate (TP*_i_*_→_*_i_*), because the corresponding cross-state TP is fully determined by 1 − TP*_i_*_→_*_i_* and is therefore mathematically redundant (i.e., TP*_i_*_→_*_j_* = 1 − TP*_i_*_→_*_i_*).

To test whether mazindol altered the balance between metastates, we fitted beta generalized linear mixed-effects models (GLMM) with a logit link. FO and TP are continuous proportions bounded between 0 and 1; beta regression respects this bounded support and naturally accommodates the mean–variance relationship typical of proportion data [84], while the mixed-effects structure accounts for within-participant dependence. The fixed-effects structure included drug condition, metastate, and their interaction. Random effects included a participant-specific random intercept and a random slope for the metastate factor. Inference on the interaction term was based on a likelihood-ratio test comparing models with and without the interaction term, accommodating potential boundary issues in variance estimates [85]. Importantly, we interpret the interaction term as a direct test of a mazindol-induced shift in *metastate balance*, that is, whether the relative difference between metastates depends on drug condition.

For both FO and TP outcomes, the interaction term in the GLMM constituted the primary inferential test. As a prespecified secondary analysis to aid interpretation of the direction of effects underlying the interaction, we evaluated the planned simple-effect contrasts with two-sided sign-flip permutation paired *t*-tests. For FO, the planned contrasts were the between-metastate differences under placebo and under drug. For TP, because the outcome was the stay probability TP_*i*→*i*_ for each source metastate, the planned contrasts were the drug–placebo differences calculated separately for each source metastate.

## 3. Results

### 3.1 PET evidence for dopamine release

One-sample t-tests on percent decreases in striatal *BP*_ND_ showed significant changes for both tracers ([^11^C]raclopride: *t_15_* = 2.637, *p* = 0.019, Cohen’s *d* = 0.659; [^18^F]FE-PE2I: *t_15_* = 18.716, *p* = 8.231 × 10^−12^, *d* = 4.679). These effects indicate that mazindol occupied presynaptic DAT sites and elevated endogenous extracellular dopamine concentration in the striatum (Fig. 3A).

**Figure 3.**
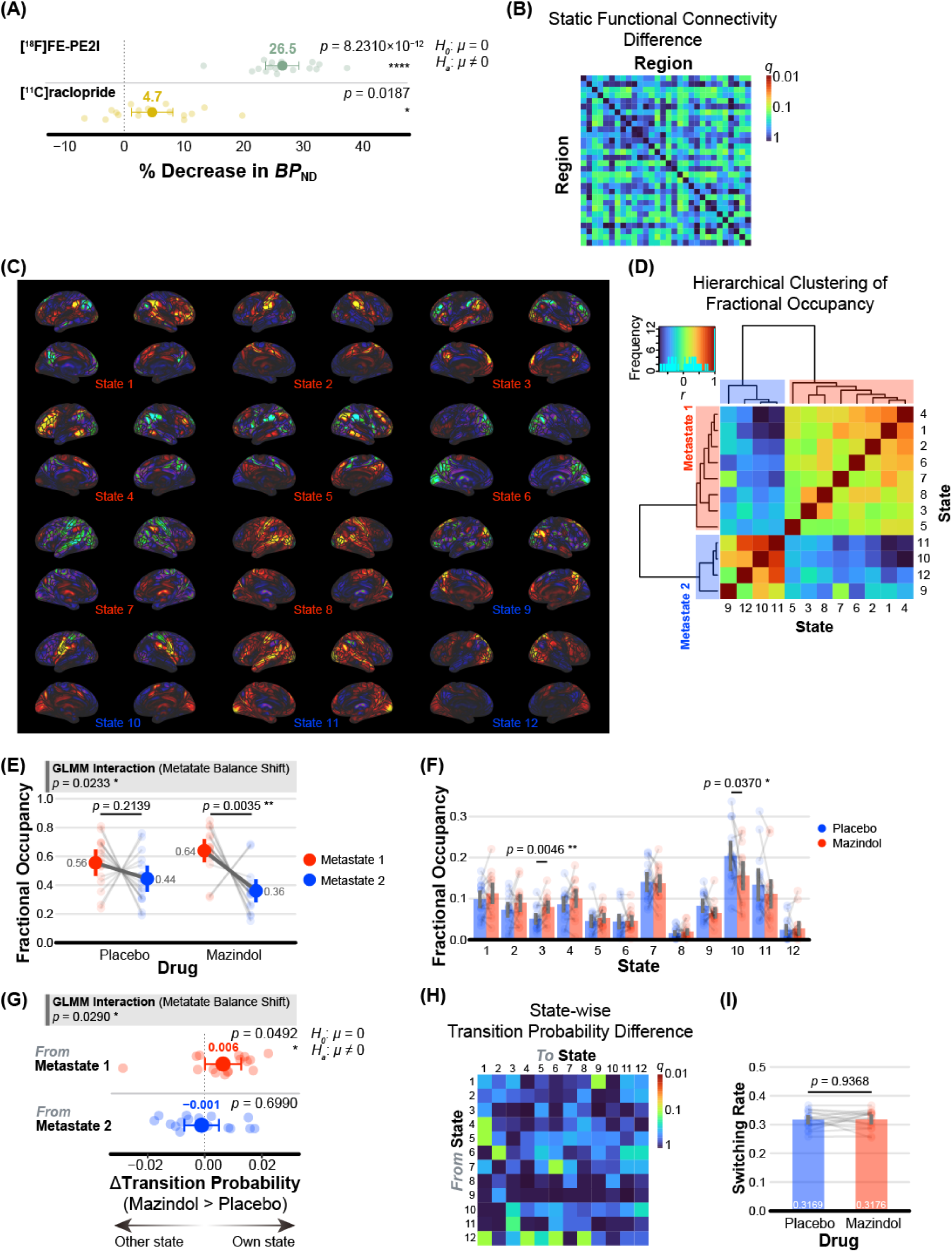
PET and fMRI results. (A) Tracer-wise percent decrease in *BP*_ND_. Dual-tracer PET indicates DAT blockade and increased extracellular dopamine following mazindol administration. Small dots indicate individual data points, and large dots indicate the mean. (B) Conventional sFC results for DMN-related 30 parcels. Colors represent between-condition differences. No difference was observed between the mazindol and placebo conditions (*q >* 0.05). (C) Twelve HMM state maps. Metastate 1 (states 1–8) anchored in higher-order cognitive regions, including the DMN, language, and salience networks, whereas metastate 2 (states 9–12) predominantly encompasses sensorimotor and perceptual cortices, including visual and auditory cortices. (D) Two distinct metastates confirmed by hierarchical clustering of the fractional occupancy (FO) correlation matrix. Correlation coefficients are color-coded using the color key shown at the top-left histogram. (E) FO for metastates. Transparent dots and lines represent individual data, and large dots indicate the mean. (F) State-wise FO. Transparent dots and lines represent individual data, and bars show the mean. (G) Transition probability (TP) difference (mazindol > placebo) from metastates. Data are shown as in A. (H) State-wise TP. Colors represent between-condition differences. No difference was observed between the mazindol and placebo conditions (*q >* 0.05). (I) Switching rate between states. No difference was observed between the mazindol and placebo conditions. Data are shown as in F. Error bars indicate the 95% CI of the mean.

### 3.2 Static functional connectivity

Using the 148 × 148 parcel-wise correlation matrices derived from rfMRI data, we compared the mazindol and placebo conditions with paired t-tests on Fisher Z-transformed coefficients. We found no significant differences in sFC between the conditions after FDR correction (*q* > 0.05). Planned confirmatory analysis of DMN-related 30 × 30 parcel-wise correlation matrices also did not detect any significant sFC differences between conditions (Fig. 3B; *q* > 0.05).

### 3.3 Brain states and metastates

HMM analysis of rfMRI identified twelve recurrent brain states, broadly recapitulating the canonical organization described by Vidaurre et al. [5] (Fig. 3C). Modularity-maximization-based clustering of the TP matrix and hierarchical clustering of the FO correlation matrix (Fig. 3D) both converged on two higher-order metastates, each nesting multiple states (states 1-8 for metastate 1, states 9-12 for metastate 2), demonstrating a hierarchical organization aligned with this prior framework. As in Vidaurre et al. [5], one metastate clustered associative circuitry supporting higher cognition, including the DMN, language, and salience networks, whereas the other comprised sensorimotor and perceptual regions, primarily visual and auditory cortices.

The FO beta GLMM showed a significant condition × metastate interaction (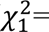= 5.145, *p* = 0.023), indicating a drug-induced shift in metastate balance (Fig. 3E). Planned contrasts revealed a significant effect of mazindol on FO. Under placebo, FO did not differ between metastates (*p* = 0.214, *d* = 0.322), whereas under mazindol, FO was significantly higher for metastate 1 than for metastate 2 (*p* = 0.004, *d* = 0.923). Furthermore, we compared state-wise FO between the mazindol and placebo conditions using paired t-tests (Fig. 3F). We found a significant prolongation for state 3 (DMN, *t_15_* = 3.323, *p* = 0.005, *d* = 0.831) and a significant shortening for state 10 (somatosensory network, *t_15_* = 2.289, *p* = 0.037, *d* = 0.572) in the mazindol relative to the placebo condition (all other states, *p* > 0.05). The difference in state 3 survived FDR correction (*q* = 0.037), but that in state 10 did not (*q* = 0.148). These results indicate that, compared with placebo, mazindol selectively increased time spent in the DMN state and metastate 1, while decreasing time spent in metastate 2 overall.

Analyses of TP also confirmed the significant lengthening of metastate 1 (Fig. 3G). The FO beta GLMM showed a significant condition × metastate interaction (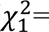= 4.767, *p* = 0.029). Planned analysis showed a significant increase in the own-state TP for metastate 1 and a corresponding significant decrease in cross-state TP from metastate 1 to 2 (*p* = 0.049, *d* = 0.537), confirming that participants spent more time in metastate 1. We observed no significant change in the own-state TP for metastate 2 or in the cross-state TP from metastate 2 to 1 (*p* = 0.699, *d* = 0.097). Then, we compared state-wise transition probabilities between the mazindol and placebo conditions (Fig. 3H) and found no significant individual TPs (*q* > 0.05). Together, the results indicate that mazindol shifts brain metastate dynamics largely by regulating transitions originating from metastate 1, with effects emerging more clearly when states are considered at the metastate level.

In addition, we evaluated the between-state switching rate using a paired t-test (Fig. 3I). We observed no significant difference between mazindol and placebo conditions in switching rate (*t_15_* = 0.081, *p* = 0.937, *d* = 0.020). Thus, we observed a redistribution of occupancy favoring metastate 1, particularly the DMN, manifested as an increased tendency to remain in metastate 1 (mathematically equivalent to a decelerated transition from metastate 1 to 2), while the overall switching rate remained unchanged.

### 3.4 Correlation between extracellular dopamine and brain states

Figure 4 shows positive correlations between the percent decreases in striatal *BP*_ND_ for [^11^C]raclopride and the ΔFO (mazindol > placebo) of both state 4 (executive network, *r* = 0.518, *p* = 0.040) and state 6 (prefrontal and reduced visual network, *r* = 0.604, *p* = 0.013). Both associations were corroborated by Kendall’s *τ* (all *p* < 0.05) but did not survive FDR correction (all *q* > 0.05) and should be interpreted cautiously. Correlations with other states were small and non-significant (all *p* > 0.05). Although exploratory, the observed directions are consistent with our hypothesis that striatal dopamine release is associated with the prolonged occupancy of particular cognitive brain states.

**Figure 4.**
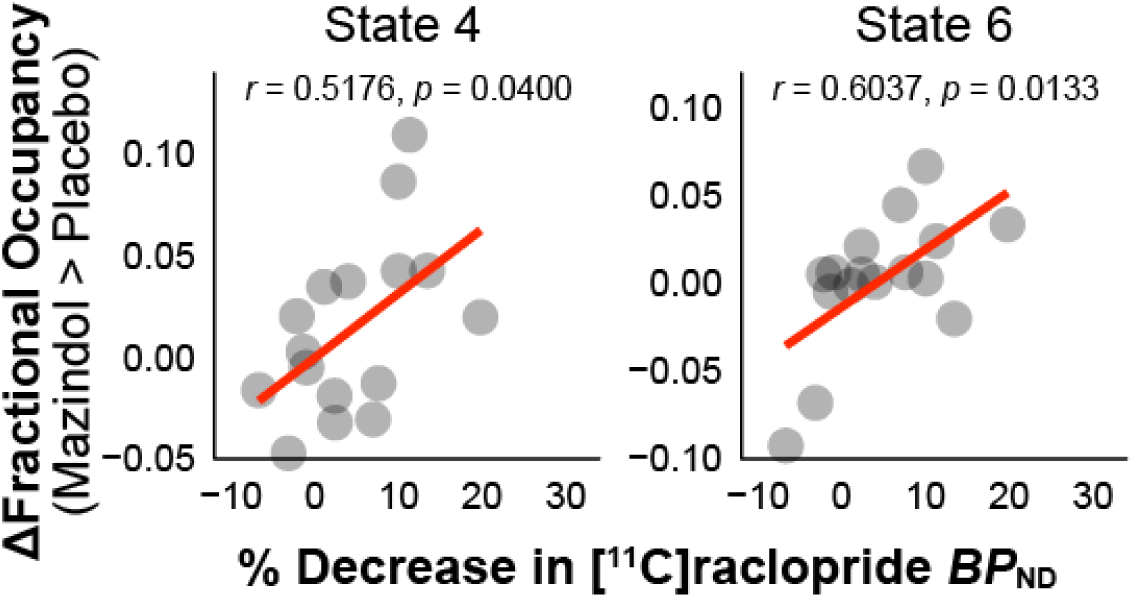
Correlation between striatum dopamine and cognitive brain states’ FO (mazindol > placebo). Dots represent individual data, and lines represent linear regression lines using ordinary least squares.

In a supplementary analysis, the percent decrease in striatal [^18^F]FE-PE2I *BP*_ND_ was correlated with the ΔFO of state 5 (*r* = −0.596, *p* = 0.015); however, this association appeared to be driven by one influential data point and was not supported by Kendall’s *τ* (*p* > 0.05).

## 4. Discussion

We found that pharmacological modulation with mazindol was associated with systematic changes in large-scale brain dynamics. Rather than yielding a simple increase or decrease in connectivity, mazindol shifted the balance between two distinct, functionally opposing brain metastates: it prolonged the time spent in a cognitive metastate, characterized by a prominent DMN state, while reducing the duration of a sensorimotor-perceptual metastate. Exploratory correlations with striatal dopamine release measured by [^11^C]raclopride PET were consistent with the drug effects (i.e., a positive correlation between dopamine release and cognitive metastate occupancy), although they did not survive correction for multiple comparisons. Taken together, these findings suggest that catecholaminergic modulation is closely linked to the temporal dynamics of brain states and may help reconcile apparently conflicting findings on dopamine’s influence on the DMN.

This temporal stabilization of the cognitive metastate offers a macroscopic account of dopamine’s established functions in cognition. At the circuit level, dopamine is thought to stabilize neural representations and increase the signal-to-noise ratio of neuronal firing, thereby prioritizing relevant activity over background activity [17, 19, 22]. Our results extend this principle to the network level: by prolonging the DMN-prominent cognitive metastate and shortening the sensorimotor-perceptual metastate, catecholaminergic modulation appears to bias large-scale dynamics toward more stable, sustained internal cognition while attenuating competing exogenous sensory inputs. This dynamic perspective also clarifies why conventional sFC measures have yielded mixed results. Because sFC reduces an fMRI time series to a single time-averaged estimate, it is insensitive to drug-induced shifts in how time is allocated across states, such that the same underlying reallocation can manifest as an increase, a decrease, or no net change in connectivity. By contrast, our HMM-based dFC approach explicitly models time-varying brain states and their dynamics, making it more sensitive to such effects. Accordingly, time-resolved connectivity may be a key feature for uncovering how neuromodulatory systems shape brain function.

Our findings also suggest a direct neurochemical influence on the dynamic balance along the brain’s principal gradient. This gradient runs from unimodal sensorimotor and perceptual to transmodal association cortices, with the DMN at its apex [28, 47, 48], consistent with a functional opponency between externally oriented processing and internally oriented cognition [5, 86]. By showing that mazindol biases the brain’s temporal landscape toward a DMN-led cognitive metastate, our results suggest that catecholaminergic tone promotes internally oriented cognitive processing over externally driven sensory processing.

This dynamic reframing may also offer new insights into the pathophysiology of dopamine-related disorders. In conditions like ADHD, DMN dysregulation is linked to intrusive, spontaneous mind-wandering and an inability to suppress the DMN during demanding tasks [87–89]. In Parkinson’s disease, cognitive deficits are associated with altered DMN connectivity and a pathological coupling with the executive control network [90–92]. The therapeutic efficacy of dopaminergic medications in these conditions, which has been linked to normalization of DMN function [57–60], may reflect not a simple increase or decrease in DMN connectivity but a restoration of normative temporal stability in large-scale brain dynamics.

Our findings also complement and extend prior work by Shine et al. [25], who used time-windowed graph-theoretical analyses in Parkinson’s disease to show that dopaminergic state reshapes the balance between segregated and integrated network configurations. Whereas they described how dopamine depletion altered global topology along a segregated–integrated continuum, our HMM-based approach provides a discrete, state-level decomposition of these dynamics in the healthy brain. Rather than focusing solely on a global segregation–integration axis, we characterize dopaminergic influences in terms of how the brain allocates time across metastates that contrast sensorimotor/perceptual systems with cognitive systems. Although the two analytic frameworks are not directly commensurable, both sets of results are consistent with the view that dopaminergic tone regulates the repertoire and temporal profile of large-scale network configurations, extending beyond what can be inferred from conventional time-averaged sFC measures. By further combining fMRI with [¹¹C]raclopride-PET, our exploratory analyses raise the possibility that inter-individual variation in striatal dopamine release is associated with the expression and temporal dynamics of these brain states, offering a neurochemically informed perspective on how dopamine modulates large-scale brain-state dynamics.

One limitation of our study is that the pharmacological agent used, mazindol, is a potent inhibitor of both the DAT and the noradrenaline transporter (NAT). The known impacts of mazindol on noradrenaline suggest a potential contribution from this system as well. Given that the noradrenaline system is a potent modulator of arousal and network state [93, 94], understanding the interplay between these two catecholamine systems is a critical next step [44]. Future research could aim to disentangle the respective contributions of dopamine and noradrenaline to the regulation of brain-state dynamics, clarifying how these systems may work in concert to produce the effects observed here.

Second, because our PET measurements relied on [^11^C]raclopride, a D2 receptor radioligand sensitive to changes in striatal dopamine, our findings primarily reflect D2 receptor-mediated signaling. However, dopamine’s cognitive effects are also critically mediated by mesocortical pathways and extrastriatal D1 and D2 receptors, which may exert a more direct impact on DMN-linked metastate dynamics. The dynamic state stabilization observed here is thus likely to arise from a complex interplay between striatal and extrastriatal mechanisms, including D1- and D2-mediated maintenance of cognitive representations [23, 95, 96]. Future research combining dynamic fMRI with extrastriatal dopamine-sensitive PET ligands targeting prefrontal and DMN regions will be essential to dissect this relationship and test more directly how dopamine shapes DMN dynamics.

Although this study examined dopamine-modulated dynamics at rest, an important question is how they generalize to periods of active task engagement. Resting-state activity is not entirely divorced from task-related activity and cognition: prestimulus DMN activity can predict subsequent behavioral performance and task-evoked brain activity [97, 98], suggesting that the baseline dynamics we measured are directly relevant for cognitive engagement. Future studies should clarify how dopaminergic modulation of these baseline brain-state dynamics constrains the reconfiguration of large-scale networks and performance during active task engagement.

Furthermore, both our exploratory PET findings and prior work highlight the importance of individual differences. Baseline neurochemistry appears to shape how the brain’s dynamic organization responds to pharmacological challenge. This principle may extend to higher-order traits or clinical symptoms. For instance, impulsivity, which has been linked to baseline striatal dopamine function [99], predicts frontostriatal and working-memory responses to a dopamine D2 receptor agonist [100]. High-impulsive (putatively low D2) individuals showed cognitive improvement with the drug, whereas low-impulsive individuals did not. This pattern suggests that the dopamine-induced stabilization of the cognitive metastate may be most pronounced, and potentially most beneficial, in individuals with specific traits or conditions. Future research integrating trait or symptom measures, neuroimaging, and pharmacology is crucial for developing a more personalized understanding of neuromodulation.

In conclusion, the present study helps reconcile the apparently paradoxical findings regarding dopamine’s role in regulating the DMN. By shifting the analytical focus from static snapshots to time-varying connectivity, we show that catecholaminergic modulation is associated with a redistribution of time across brain states that stabilizes the DMN-led cognitive metastate relative to the sensorimotor-perceptual metastate. This work highlights the importance of employing time-resolved, dynamic analytical approaches (e.g., dFC) for accurately characterizing the influence of neuromodulatory systems on brain function. By linking molecular-level neurochemistry to the macroscopic, temporal organization of brain states, our findings offer an integrative framework for understanding brain function and dysfunction. From this perspective, the pathophysiology of disorders involving disrupted neurotransmitter systems may be defined not only by static patterns of network hyper- or hypo-connectivity, but also by abnormalities in network dynamics—a failure to appropriately enter, sustain, or transition between functionally relevant brain states. Such a mechanistic reframing may offer a useful target for therapeutic interventions, shifting the goal from simply modulating network activity to restoring the temporal integrity of large-scale brain dynamics.

## Acknowledgements

This study was supported by grants from the JST Moonshot R&D Grant (JPMJMS2295 to MY).

## Author contributions

YK and MY designed the study. YK, KT, YI, CS, MZ, MH, TS, and MY contributed to data collection and PET tracer–related materials. DM analyzed the data. DM and MY interpreted the results. DM wrote the initial draft. MH, TS, and MY supervised the entire project. All authors have reviewed and approved the final manuscript.

